## Supplementary material for "Towards Parsimonious Generative Modeling of RNA Families": SI

October 19, 2023

#### 1 eaDCA: Details and Analytical Derivations

##### 1.1 Analytical Derivations

In this section, we provide the analytical derivation of the iterative model updates. At each iteration, we aim at improving the generative model

$$
\begin{aligned}
P_t(a_1,\dots,a_L) &= \frac{1}{Z_t}\exp\big\{-E_t(a_1,\dots,a_L)\big\} \\
-E_t(a_1,\dots,a_L) &= \sum_{i=1}^{L} h_i(a_i) + \sum_{(ij)\in\mathcal{E}_t} J_{ij}(a_i,a_j)
\end{aligned}
\tag{1}
$$

by changing a single coupling matrix between a single pair of positions, with the aim of maximizing the model log-likelihood given the training data $\mathcal{D} = (a_i^r \,|\, i=1,\dots,L;\, r=1,\dots,M)$. Here, $\mathcal{E}_t$ is the set of currently "activated" couplings, i.e. the set of all position pairs currently connected by a non-zero coupling. The log-likelihood can be written as

$$
\log\mathcal{L}_t = \sum_{r=1}^{M} \omega_r \log P_t(a_1^r,\dots,a_L^r), \tag{2}
$$

with $\omega_r$ being the standard sequence reweighting explained in the main text, and usually applied in DCA. Using Eq. (1) and the definition $M_{\text{eff}} = \sum_r \omega_r$ of the effective sequence number, we have

$$
\log\mathcal{L}_t = -M_{\text{eff}}\log Z_t + \sum_{r=1}^{M}\omega_r\left(\sum_{i=1}^{L} h_i(a_i^r) + \sum_{ij\in\mathcal{E}_t} J_{ij}(a_i^r,a_j^r)\right) . \tag{3}
$$

To obtain the model $P_{t+1}(a_1,\dots,a_L)$, we add a $\Delta J_{mn}(a,b)$ to the couplings of $-E_t(a_1,\dots,a_L)$ in Eq. (1); this is done for a single position pair $1 \le m < n \le L$, but arbitrary entries $a,b \in \{A,C,G,U,-\}$:

$$
\log\mathcal{L}_{t+1} = -M_{\text{eff}}\log Z_{t+1} + \sum_{r=1}^{M}\omega_r\left(\sum_{i=1}^{L} h_i(a_i^r) + \sum_{ij\in\mathcal{E}_t} J_{ij}(a_i^r,a_j^r) + \Delta J_{mn}(a_m^r,a_n^r)\right) \tag{4}
$$

We aim at finding the $\Delta J_{mn}$ maximizing the likelihood gain

$$
\Delta\log\mathcal{L} = \log\mathcal{L}_{t+1} - \log\mathcal{L}_t = -M_{\text{eff}}\log\frac{Z_{t+1}}{Z_t} + \sum_{r=1}^{M}\omega_r\Delta J_{mn}(a_m^r,a_n^r) . \tag{5}
$$

This can be simplified using the empirical two-point frequencies $f_{mn}(a,b) = \frac{1}{M_{\text{eff}}}\sum_r \omega_r \delta_{a,a_m^r}\delta_{b,a_n^r}$:

$$
\frac{\Delta\log\mathcal{L}}{M_{\text{eff}}} = -\log\frac{Z_{t+1}}{Z_t} + \sum_{a,b}\Delta J_{mn}(a,b) f_{mn}(a,b) . \tag{6}
$$

The ratio $\frac{Z_{t+1}}{Z_t}$ of the partition functions can be expressed as

$$\frac{Z_{t+1}}{Z_t} = \frac{\sum_{a_1,a_2,\dots,a_L} \exp\{-E_t(a_1,\dots,a_L)\} \cdot \exp\{\Delta J_{mn}(a_m,a_n)\}}{\sum_{a_1,a_2,\dots,a_L} \exp\{-E_t(a_1,a_2,\dots,a_L)\}} , \tag{7}$$

resulting in a simple mean value over $P_t$

$$\frac{Z_{t+1}}{Z_t} = \left\langle e^{\Delta J_{mn}(a_m,a_n)} \right\rangle_{P_t} = \sum_{a,b} e^{\Delta J_{mn}(a,b)} P^t_{mn}(a,b) , \tag{8}$$

with $P^t_{mn}(a,b)$ being the two-site marginal of model $P_t$ for positions $m,n$. Substituting the last expression back into Eq. (6), we can express the likelihood change explicitly in terms of the changed coupling and the marginal probability of the old model at iteration $t$:

$$\frac{\Delta \log \mathcal{L}}{M_{\text{eff}}} = -\log\left(\sum_{a,b} e^{\Delta J_{mn}(a,b)} P^t_{mn}(a,b)\right) + \sum_{a,b} \Delta J_{mn}(a,b) f_{mn}(a,b) \tag{9}$$

For any given pair $(m,n)$, we can maximize this expression over $\Delta J_{mn}(a,b)$ by solving

$$0 = f_{mn}(a,b) - \frac{e^{\Delta J^*_{mn}(a,b)} P^t_{mn}(a,b)}{\sum_{c,d} e^{\Delta J^*_{mn}(c,d)} P^t_{mn}(c,d)} , \tag{10}$$

i.e. by choosing

$$\Delta J^*_{mn}(a,b) = \log\left(\frac{f_{mn}(a,b)}{P^t_{mn}(a,b)}\right) . \tag{11}$$

For given $(m,n)$, the maximally realizable likelihood gain therefore reads

$$\frac{\Delta \log \mathcal{L}}{M_{\text{eff}}} = \sum_{a,b} f_{mn}(a,b) \log\left(\frac{f_{mn}(a,b)}{P^t_{mn}(a,b)}\right) = D_{KL}\left(f_{mn} \,||\, P^t_{mn}\right) . \tag{12}$$

To finalize the derivation of our algorithm we need to select, out of all possible position pairs, the one realizing the largest likelihood gain,

$$(kl) = \underset{1\le m<n\le L}{\text{argmax}}\; D_{KL}\left(f_{mn} \,||\, P^t_{mn}\right) , \tag{13}$$

and update only the corresponding coupling

$$\Delta J^*_{kl}(a,b) = \log\left(\frac{f_{kl}(a,b)}{P^t_{kl}(a,b)}\right) \tag{14}$$

for all $a,b \in \{A,C,G,U,-\}$. Note that Eq. (13) optimizes over all pairs of positions, including those already present in $\mathcal{E}_t$. This choice results in the following change in the Potts Model:

$$\begin{aligned} E_{t+1}(a_1,\dots,a_L) &= E_t(a_1,\dots,a_L) - \Delta J^*_{kl}(a_k,a_l) , \\ \mathcal{E}_{t+1} &= \mathcal{E}_t \cup \{(kl)\} . \end{aligned} \tag{15}$$

#### 1.2 Monte Carlo Sampling

Due to the high computational cost of calculating the exact two-point probabilities $P^t_{kl}(a,b)$ for the coupling update in Eq. (14), we resort to a Monte Carlo approach to obtain an approximation. The Monte Carlo simulation involves sampling a number of states (sequences) from $P_t(a_1,a_2,\dots,a_L)$, and using the sample two-point frequencies $P^{m_1}_{kl}(a,b)$ as an estimate for the model two-point probabilities $P^t_{kl}(a,b)$. Our chosen Monte Carlo algorithm is Gibbs sampling. The default implementation use 10000 independent chains to compute $P^{m_1}_{kl}(a,b)$. To optimize computational time, we used the Persistent Contrastive Divergence technique in our implementation. Essentially, this approach initializes the Monte Carlo runs for the sampling of $P_t(a_1,a_2,\dots,a_L)$ to samples obtained from the $P_{t-1}(a_1,a_2,\dots,a_L)$ at the previous step. Given that two consecutive models differ by just one interaction coupling, the samples from $P_{t-1}$ are already close to equilibrium for $P_t$. The default implementation performs 5 Gibbs sweeps on all the 10000 training chains. Upon selecting our final generative model, we performed an independent re-sampling starting from randomized sequences. This step ensured that the Persistent Contrastive Divergence did not trap the simulation in a locally stable state.

#### 1.3 Regularization

Suppose that in our Monte Carlo we sample a sequence with a pair of nucleotides at sites $(k, l)$ that is absent from the natural dataset. In this case, if the eaDCA traverses the edge in question, the resulting coupling update Eq. (14) is $-\infty$. Similarly, if a sampled pair present in the natural dataset is never sampled, we encounter a $+\infty$ coupling update. These conditions, and in general excessively high valued couplings, pose threats to the performance and to the ergodicity of the models.

To mitigate these issues, we introduce a regularization for the update:

$$\Delta J^*_{kl}(a,b) = \log\left(\frac{(1-\alpha)f_{kl}(a,b) + \frac{\alpha}{q^2}}{(1-\alpha)P^t_{kl}(a,b) + \frac{\alpha}{q^2}}\right) \tag{16}$$

As the value of $\alpha$ increases, the regularization effect on the update correspondingly intensifies. In our models, we used a fixed $\alpha = 0.1$. This choice allowed our model to pass the ergodicity tests in the re-sampling and achieve high performance scores.

Interestingly, despite this modification, we note from Eq. (19) that the termination condition remains the same – the equality between the natural two-point natural frequencies $f_{kl}(a, b)$ and the model two-point probabilities $P_{kl}(a, b)$.

#### 1.4 Partition Function Conservation

In this section we provide the proof that eaDCA preserves the partition function $Z$. Substituting the coupling update Eq. (14) in Eq. (8) we can express the ratio $\frac{Z_{t+1}}{Z_t}$ as:

$$\frac{Z_{t+1}}{Z_t} = \sum_{a,b} \exp\left\{\log\left(\frac{f_{kl}(a,b)}{P^t_{kl}(a,b)}\right)\right\} P^t_{kl}(a,b) = \sum_{a,b} \frac{f_{kl}(a,b)}{P^t_{kl}(a,b)} P^t_{kl}(a,b) = \sum_{a,b} f_{kl}(a,b) = 1 \ , \tag{17}$$

which proves that $Z_{t+1} = Z_t$ . Since our procedure starts from the Profile Model, which has a $Z_0 = 1$, all the models in the eaDCA chain are normalized. A nice consequence is that the models entropy are easily accessible trough the relation

$$S_t =< E_t >_{P_t} \tag{18}$$

In Section 1.2 and Section 1.3 we discussed the impossibility of applying the ideal algorithm. This means that the real coupling update deviates from Eq. (14). Its real value is:

$$\Delta J^*_{kl}(a,b) = \log\left(\frac{(1-\alpha)f_{kl}(a,b) + \frac{\alpha}{q^2}}{1-\alpha)P^{m_1}_{kl}(a,b) + \frac{\alpha}{q^2}}\right) \tag{19}$$

which takes in account both the regularization term $\alpha$ and the Monte Carlo estimation $P^{m_1}_{kl}(a, b)$. This implies that Eq. (17) only holds approximately.

Using Eq. (8) and Eq. (19) we can get the real relation between $Z_{t+1}$ and $Z_t$ :

$$Z_{t+1} = Z_t \sum_{a,b} \frac{(1-\alpha)f_{kl}(a,b) + \frac{\alpha}{q^2}}{(1-\alpha)P^{m_1}_{kl}(a,b) + \frac{\alpha}{q^2}} P^t_{kl}(a,b) \ , \tag{20}$$

or

$$\log Z_{t+1} = \log Z_t + \log\left(\sum_{a,b} \frac{(1-\alpha)f_{kl}(a,b) + \frac{\alpha}{q^2}}{(1-\alpha)P^{m_1}_{kl}(a,b) + \frac{\alpha}{q^2}} P^t_{kl}(a,b)\right) . \tag{21}$$

Hence, defining

$$\Phi_t = \log\left(\sum_{a,b} \frac{(1-\alpha)f_{kl}(a,b) + \frac{\alpha}{q^2}}{(1-\alpha)P^{m_1}_{kl}(a,b) + \frac{\alpha}{q^2}} P^t_{kl}(a,b)\right) , \tag{22}$$

we can iterate Eq. (21) to obtain

$$\log Z_t = \sum_{s=0}^{t-1} \Phi_s \ . \tag{23}$$

However, we then are back to the initial problem: in order to compute $\Phi_s$ with Eq. (23), we need the exact values of $P^t_{kl}(a,b)$. A possible solution is to use another independent Monte Carlo approximation $P^{m_2}_{kl}(a,b)$ to obtain

$$\Phi_t \simeq \log\left(\sum_{a,b}\frac{(1-\alpha)f_{kl}(a,b)+\frac{\alpha}{q^2}}{(1-\alpha)P^{m_1}_{kl}(a,b)+\frac{\alpha}{q^2}}P^{m_2}_{kl}(a,b)\right) . \tag{24}$$

We tested how well these arguments hold using a very short RNA segment (first 11 residues from the RF0442 family):

| $P^{m_1}_{kl}(a,b)$ (training) sequences | $P^{m_2}_{kl}(a,b)$ $(\Phi_t)$ sequences | eaDCA iterations | Exact $S$ | Estimated $S$ |
|---|---|---|---|---|
| 1000 | 2000 | 390 | 11.56 | 11.72 |
| 1000 | 10000 | 390 | 11.65 | 11.65 |
| 1000 | 2000 | 1000 | 11.41 | 11.58 |
| 1000 | 10000 | 1000 | 11.66 | 11.66 |

**Table 1**: We illustrate the entropy estimation using 2000 and 10000 chains to compute $P^{m_2}_{kl}(a,b)$. A reduced sample of 1000 sequences, as opposed to the standard 10000, was employed for training, enabling us to examine scenarios where the violation of the conservation of $Z$ is more pronounced.

Given that this procedure results in a reasonable approximation of $\log Z_t$, we can still derive the models entropy using

$$S_t =< E_t >_{P_t} + \log Z_t \tag{25}$$

In the default eaDCA implementation we use 2000 independent chains to compute $P^{m_2}_{kl}(a,b)$.

#### 1.5 Typical eaDCA Training Process

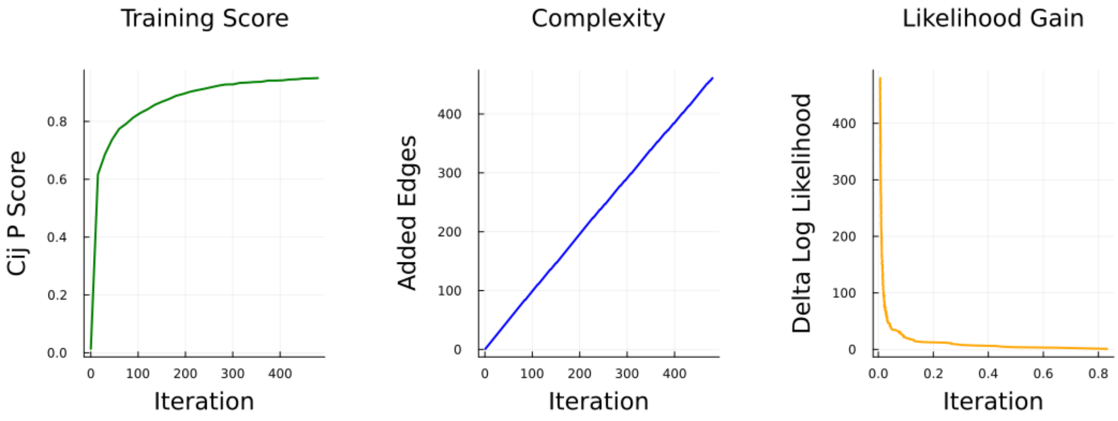


**Figure 1**: eaDCA Training Process for the RF01734 Family. The plot showcases typical behavior of the $c_{ij}$ Pearson score, the number of added edges and the likelihood gain as the number of iterations increases.

In Fig. 1 we report the typical training trajectory of eaDCA. The first few iterations yield the highest gain in Pearson Score and likelihood. After this initial spike, both the likelihood gain and the increase in score start to plateau, becoming much slower. However, delving into this slower region is essential for obtaining models with good statistical performance. Complexity, on the other hand, increases almost linearly as the number of iterations rises. While coupling-update iterations do occur, they are relatively rare, allowing us to obtain a well-performing model before they become predominant.

#### 2 Training Data And Statistical Tests

| Family Name | $L$ | $M$ | $M_{eff}$ | PR% | $S$ | $\Omega$ | PDB |
|---|---|---|---|---|---|---|---|
| RF00005 | 71 | 28770 | 2267 | 84.35% | 51.34 | $1.98 \times 10^{22}$ | 1ehz A |
| RF00028 | 251 | 611 | 233 | 88.17% | 106.31 | $1.48 \times 10^{46}$ | 1gid A Err |
| RF00050 | 140 | 4051 | 530 | 92.16% | 73.78 | $1.10 \times 10^{32}$ | 3f2q X |
| RF00059 | 105 | 12208 | 4683 | 94.34% | 77.41 | $4.16 \times 10^{33}$ | 3d2g A |
| RF00080 | 175 | 468 | 371 | 64.41% | 82.29 | $5.47 \times 10^{35}$ | 6cb3 A |
| RF00114 | 117 | 620 | 218 | 73.37% | 58.72 | $3.18 \times 10^{25}$ | 2vaz A Err |
| RF00162 | 108 | 5974 | 1283 | 91.09% | 66.23 | $5.80 \times 10^{28}$ | 3gx5 A |
| RF00167 | 102 | 2613 | 1596 | 90.18% | 71.87 | $1.63 \times 10^{31}$ | 4tzx X |
| RF00168 | 183 | 2191 | 1419 | 82.42% | 117.55 | $1.13 \times 10^{51}$ | 3dil A |
| RF00169 | 97 | 4641 | 796 | 82.50% | 64.67 | $1.22 \times 10^{28}$ | 1z43 A Err |
| RF00234 | 171 | 893 | 639 | 82.83% | 95.86 | $4.28 \times 10^{41}$ | 2h0s B Err |
| RF00379 | 136 | 3808 | 1428 | 87.83% | 89.85 | $1.05 \times 10^{39}$ | 4qln A |
| RF00380 | 169 | 1057 | 149 | 95.15% | 84.24 | $3.85 \times 10^{35}$ | 3pdr A |
| RF00442 | 108 | 845 | 312 | 89.89% | 56.02 | $2.13 \times 10^{24}$ | 5u3g B |
| RF00504 | 94 | 4395 | 1363 | 91.19% | 63.86 | $5.42 \times 10^{27}$ | 3ox0 B |
| RF01051 | 87 | 4032 | 1850 | 90.24% | 57.46 | $9.01 \times 10^{24}$ | 4yaz A |
| RF01725 | 101 | 707 | 249 | 80.32% | 57.00 | $5.69 \times 10^{24}$ | 4l81 A |
| RF01734 | 65 | 2345 | 923 | 77.84% | 44.31 | $1.75 \times 10^{19}$ | 4enc A Err |
| RF01750 | 101 | 1402 | 420 | 85.45% | 59.20 | $5.13 \times 10^{25}$ | 4xwf A |
| RF01786 | 85 | 585 | 336 | 70.39% | 48.40 | $1.05 \times 10^{21}$ | 5nwq A |
| RF01831 | 100 | 692 | 233 | 83.98% | 48.64 | $1.33 \times 10^{21}$ | 4lvv A |
| RF01852 | 91 | 2351 | 296 | 89.47% | 52.13 | $4.36 \times 10^{22}$ | 3rg5 A |
| RF01854 | 255 | 1052 | 343 | 83.15% | 92.02 | $9.20 \times 10^{39}$ | 4wfl A |
| RF02001 | 171 | 3103 | 657 | 70.21% | 96.44 | $7.64 \times 10^{41}$ | 4y1o A |
| RF02553 | 114 | 194 | 116 | 81.54% | 35.77 | $3.43 \times 10^{15}$ | 6cu1 A |

**Table 2**: Results from the application of eaDCA across all 25 test families are presented. The PDB identifiers are indicated, and if 'err' is displayed, it signifies an issue in mapping the PDB to the alignment.

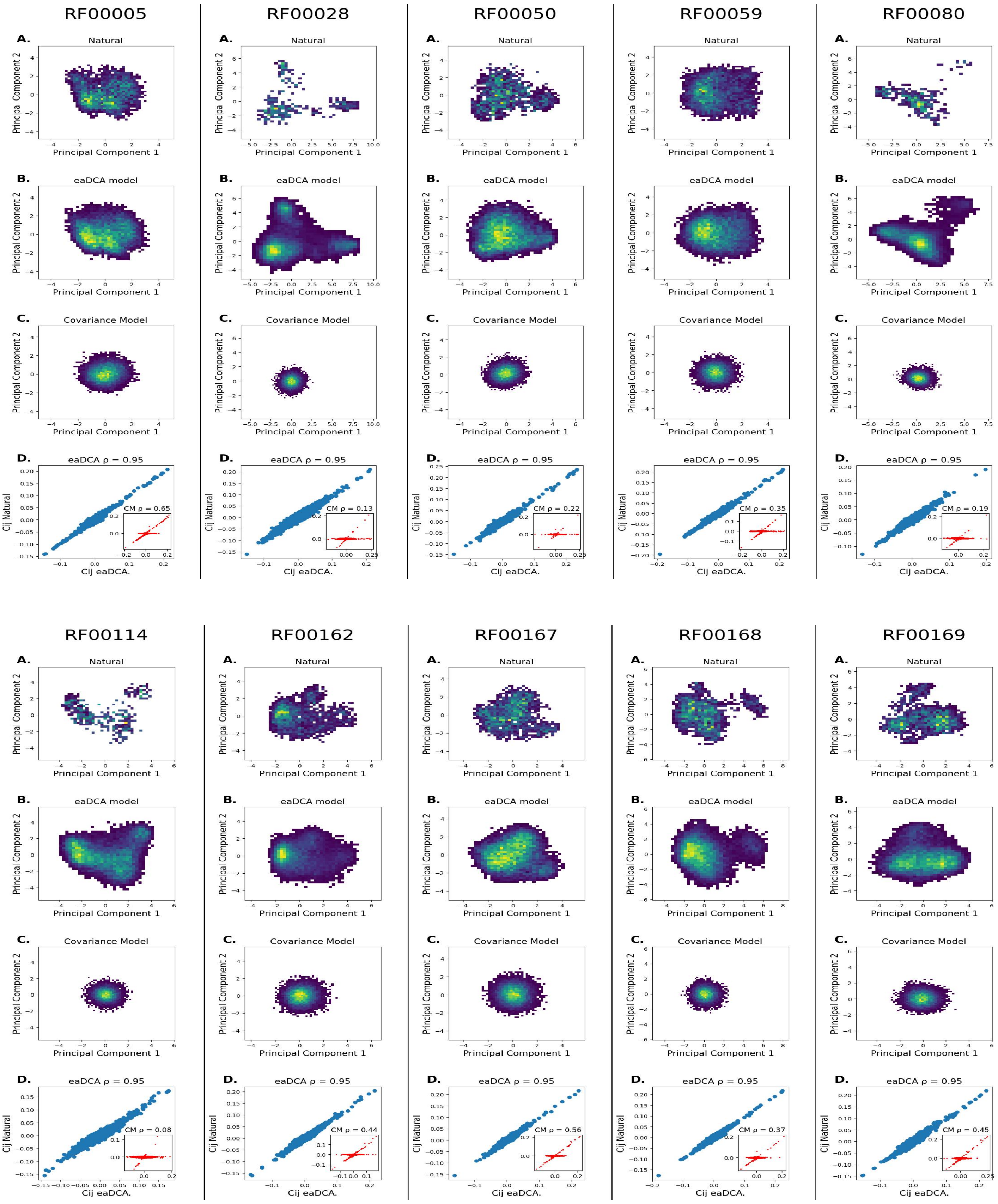
RF00005
RF00028
RF00050
RF00059
RF00080
A.
Natural
Principal Component 1
Principal Component 2
B.
eaDCA model
C.
Covariance Model
D.
eaDCA ρ = 0.95
CM ρ = 0.65
CM ρ = 0.13
CM ρ = 0.22
CM ρ = 0.35
CM ρ = 0.19
Cij Natural
Cij eaDCA.
RF00114
RF00162
RF00167
RF00168
RF00169
CM ρ = 0.08
CM ρ = 0.44
CM ρ = 0.56
CM ρ = 0.37
CM ρ = 0.45

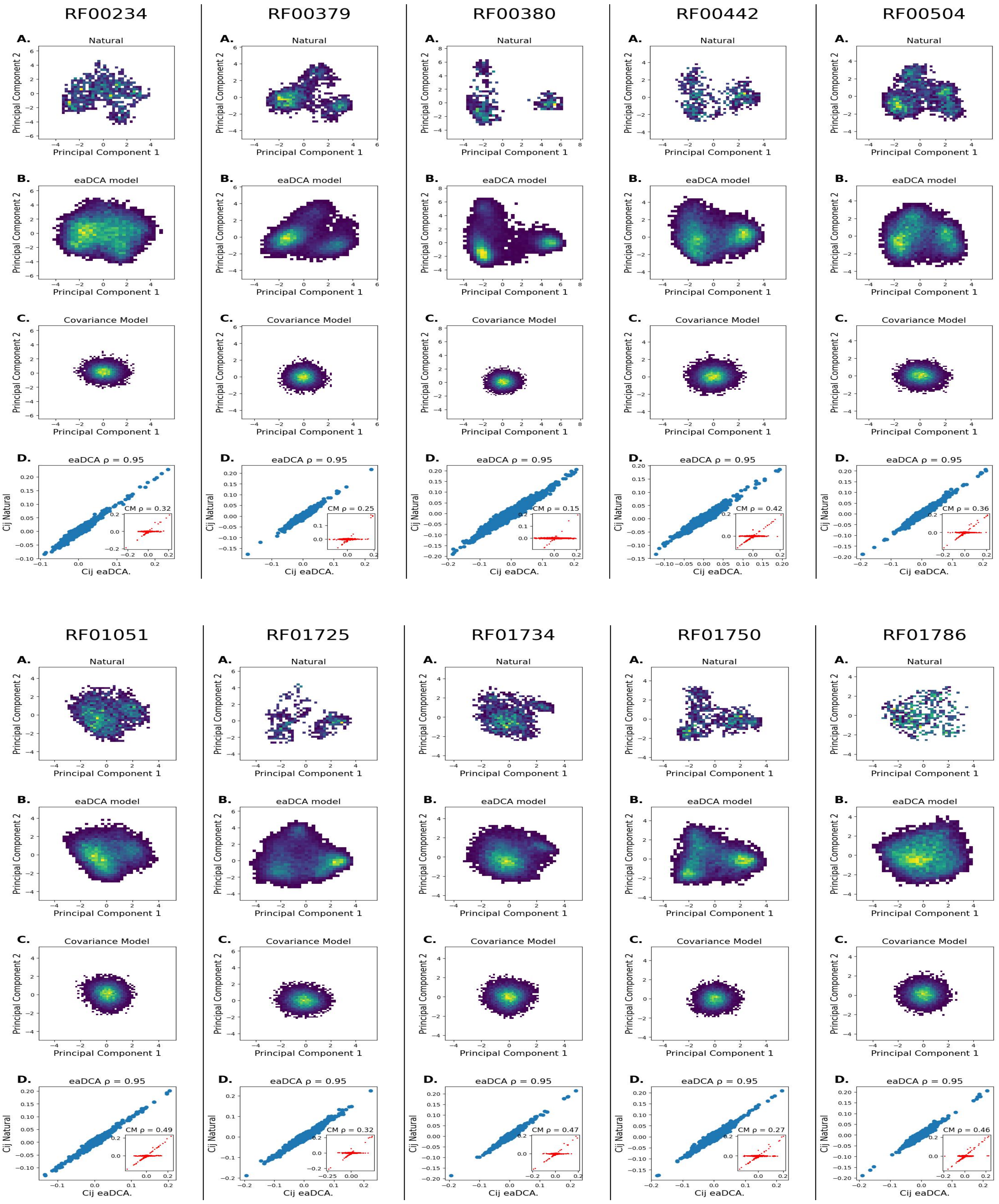
RF00234
RF00379
RF00380
RF00442
RF00504
A. Natural
B. eaDCA model
C. Covariance Model
D. eaDCA ρ = 0.95
Principal Component 1
Principal Component 2
Cij Natural
Cij eaDCA.
CM ρ = 0.32
CM ρ = 0.25
CM ρ = 0.15
CM ρ = 0.42
CM ρ = 0.36
RF01051
RF01725
RF01734
RF01750
RF01786
CM ρ = 0.49
CM ρ = 0.32
CM ρ = 0.47
CM ρ = 0.27
CM ρ = 0.46

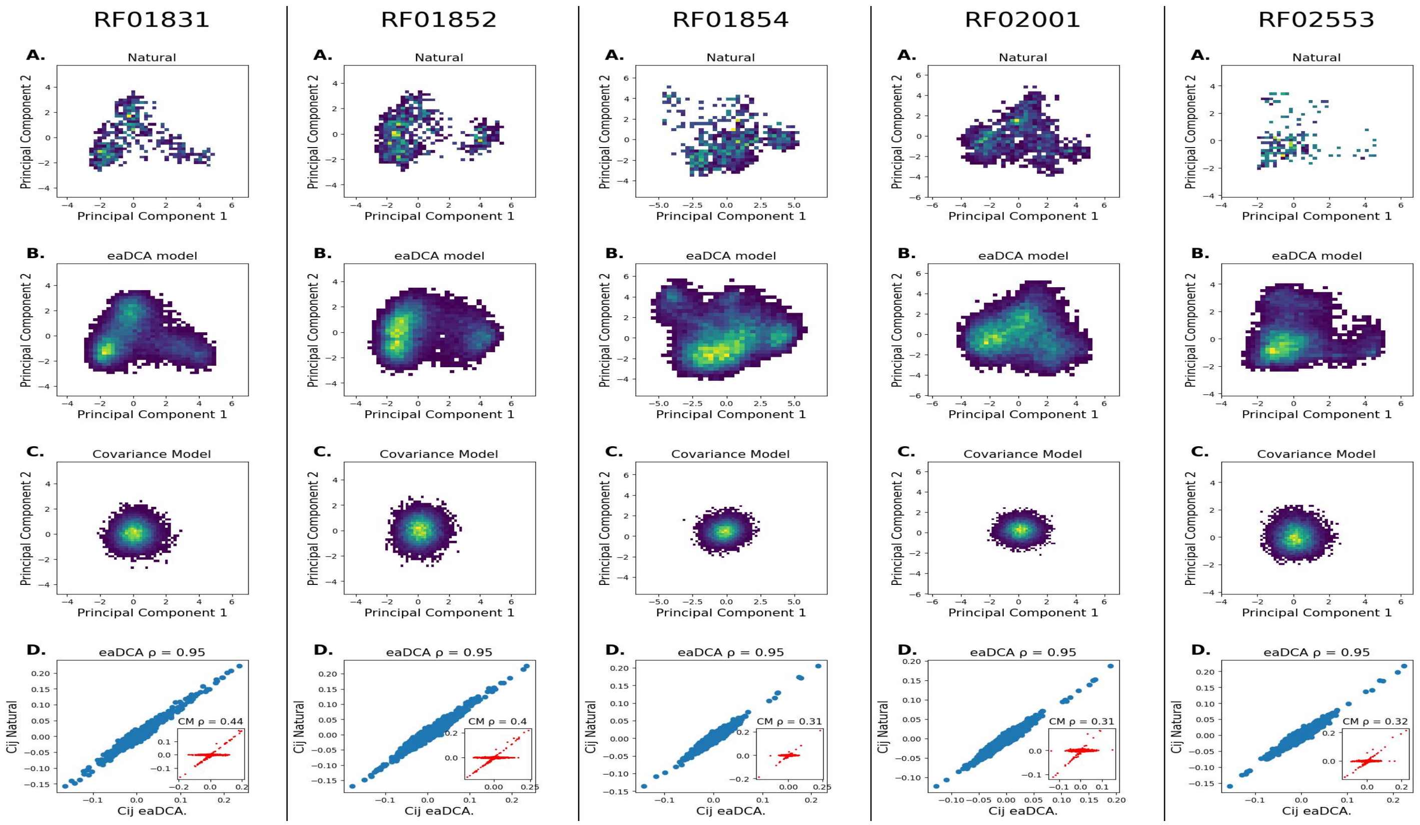


**Figure 2**: **A.** PCA dimensional reduction of natural sequences. **B.** PCA dimensional reduction of eaDCA generated sequences. **C.** PCA dimensional reduction of covariance model generated sequences. **D.** two-point statistics representation for eaDCA model and covariance model.

### 3 Parameter Interpretation

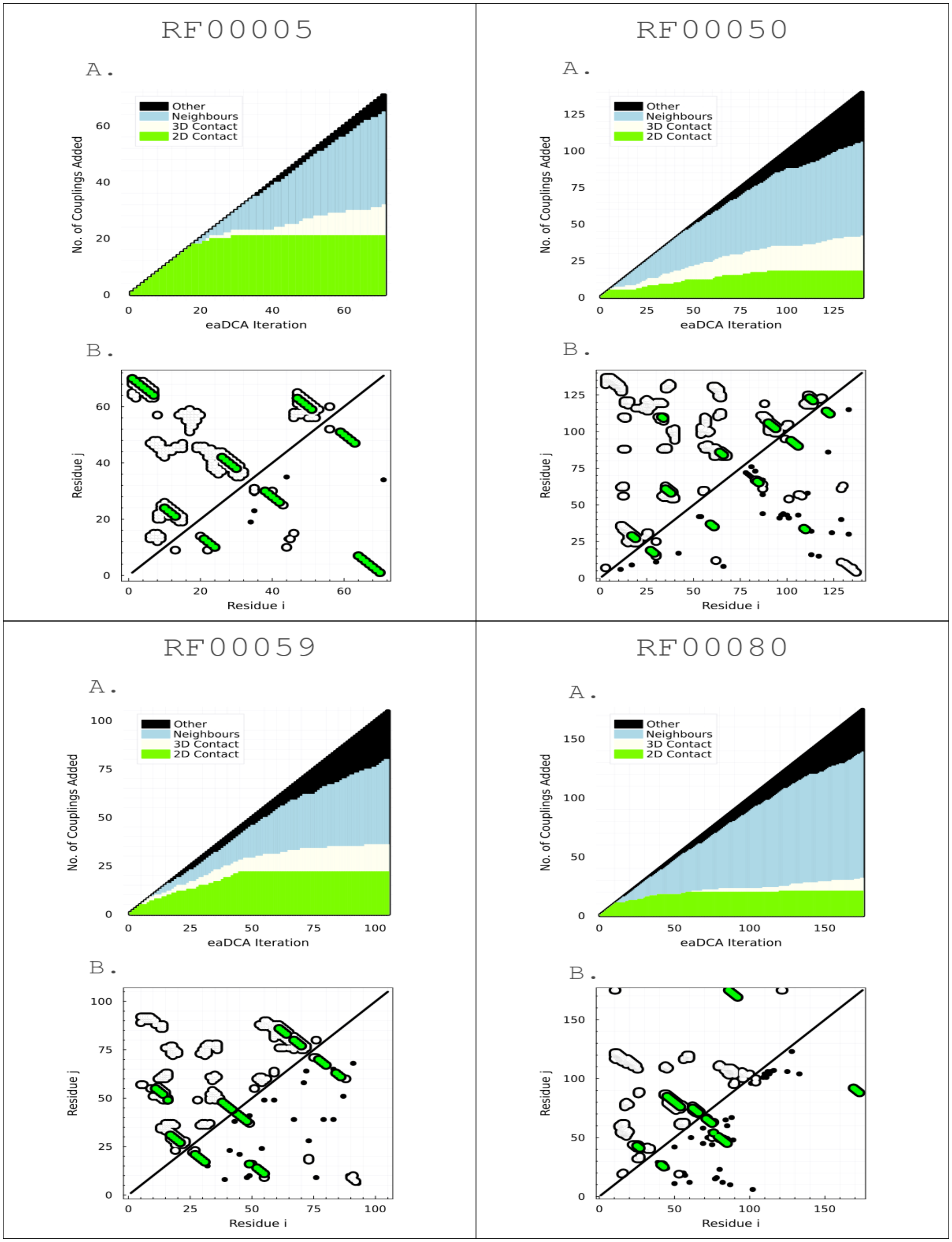

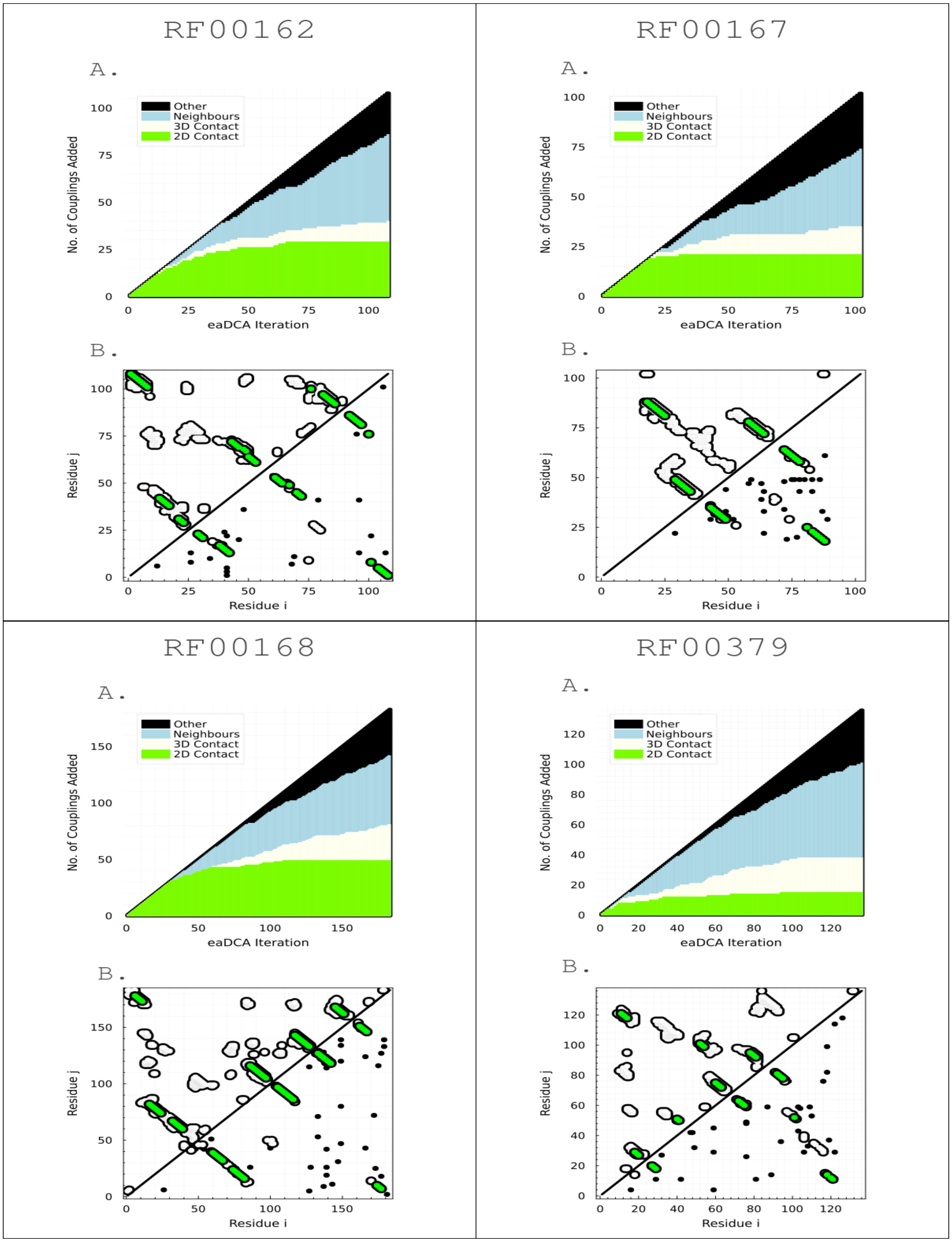
RF00162
A.
Other
Neighbours
3D Contact
2D Contact
No. of Couplings Added
eaDCA Iteration
B.
Residue j
Residue i
RF00167
A.
Other
Neighbours
3D Contact
2D Contact
No. of Couplings Added
eaDCA Iteration
B.
Residue j
Residue i
RF00168
A.
Other
Neighbours
3D Contact
2D Contact
No. of Couplings Added
eaDCA Iteration
B.
Residue j
Residue i
RF00379
A.
Other
Neighbours
3D Contact
2D Contact
No. of Couplings Added
eaDCA Iteration
B.
Residue j
Residue i

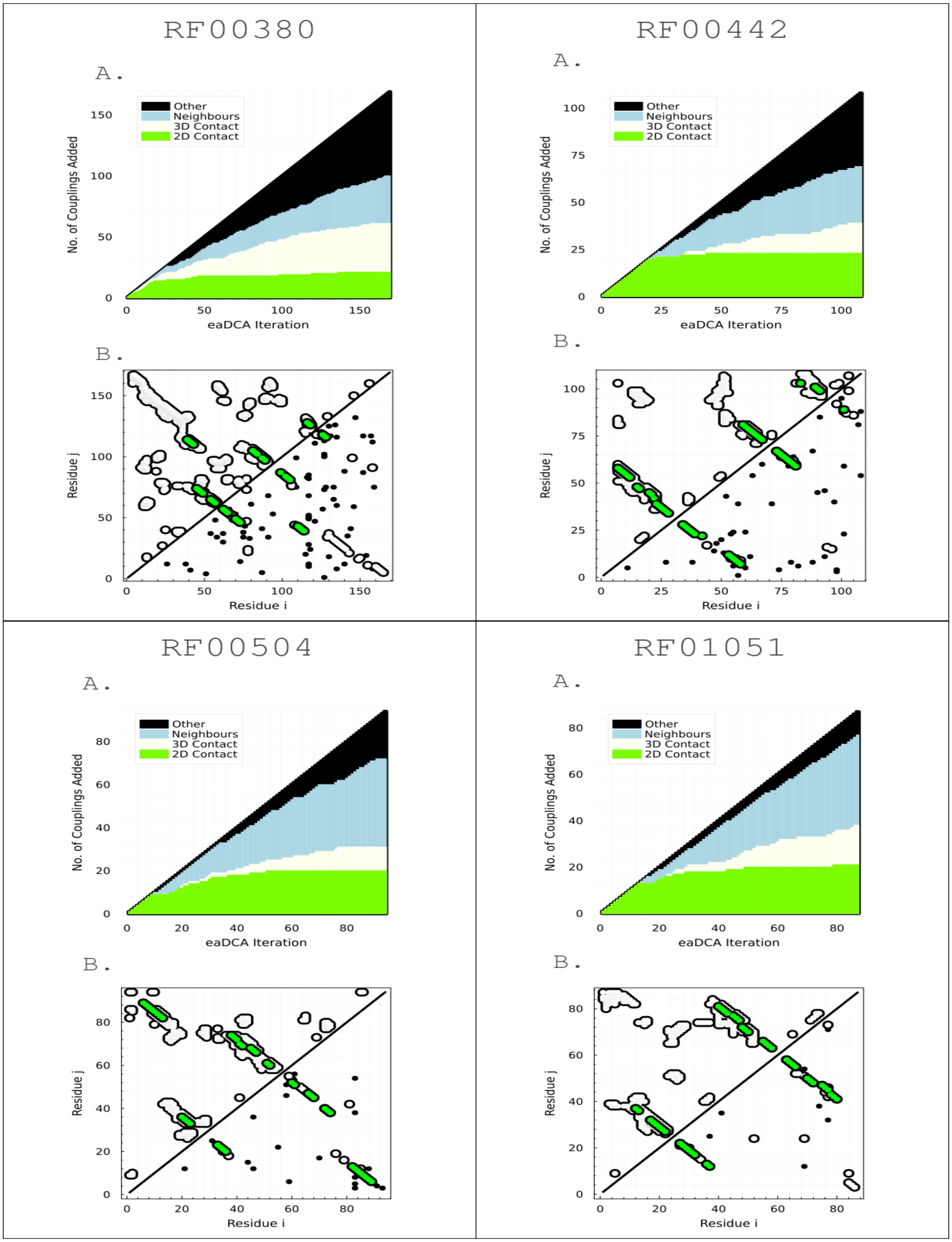
RF00380
A.
Other
Neighbours
3D Contact
2D Contact
No. of Couplings Added
eaDCA Iteration
B.
Residue j
Residue i
RF00442
A.
Other
Neighbours
3D Contact
2D Contact
No. of Couplings Added
eaDCA Iteration
B.
Residue j
Residue i
RF00504
A.
Other
Neighbours
3D Contact
2D Contact
No. of Couplings Added
eaDCA Iteration
B.
Residue j
Residue i
RF01051
A.
Other
Neighbours
3D Contact
2D Contact
No. of Couplings Added
eaDCA Iteration
B.
Residue j
Residue i

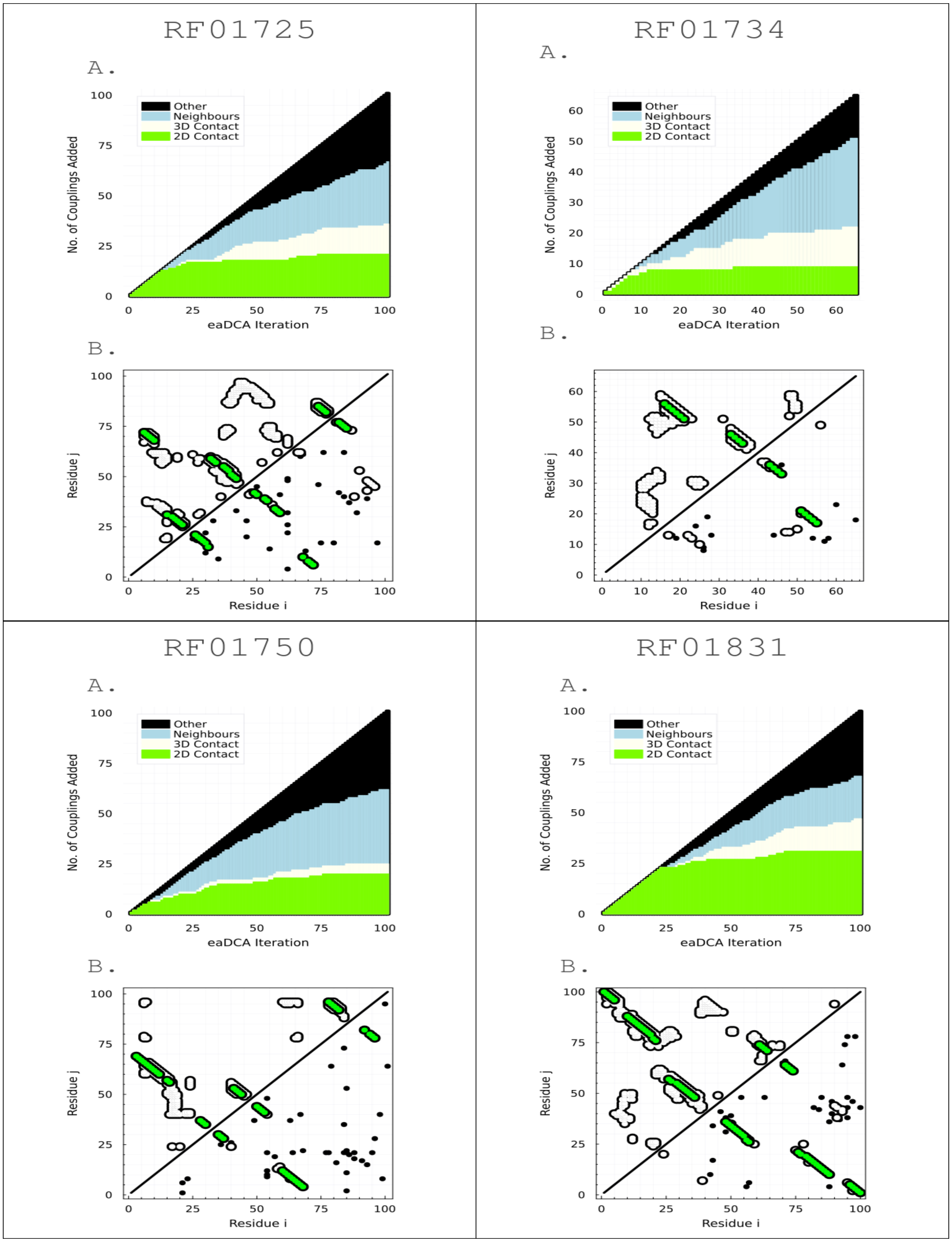
RF01725
A.
Other
Neighbours
3D Contact
2D Contact
No. of Couplings Added
eaDCA Iteration
B.
Residue j
Residue i
RF01734
A.
Other
Neighbours
3D Contact
2D Contact
No. of Couplings Added
eaDCA Iteration
B.
Residue j
Residue i
RF01750
A.
Other
Neighbours
3D Contact
2D Contact
No. of Couplings Added
eaDCA Iteration
B.
Residue j
Residue i
RF01831
A.
Other
Neighbours
3D Contact
2D Contact
No. of Couplings Added
eaDCA Iteration
B.
Residue j
Residue i

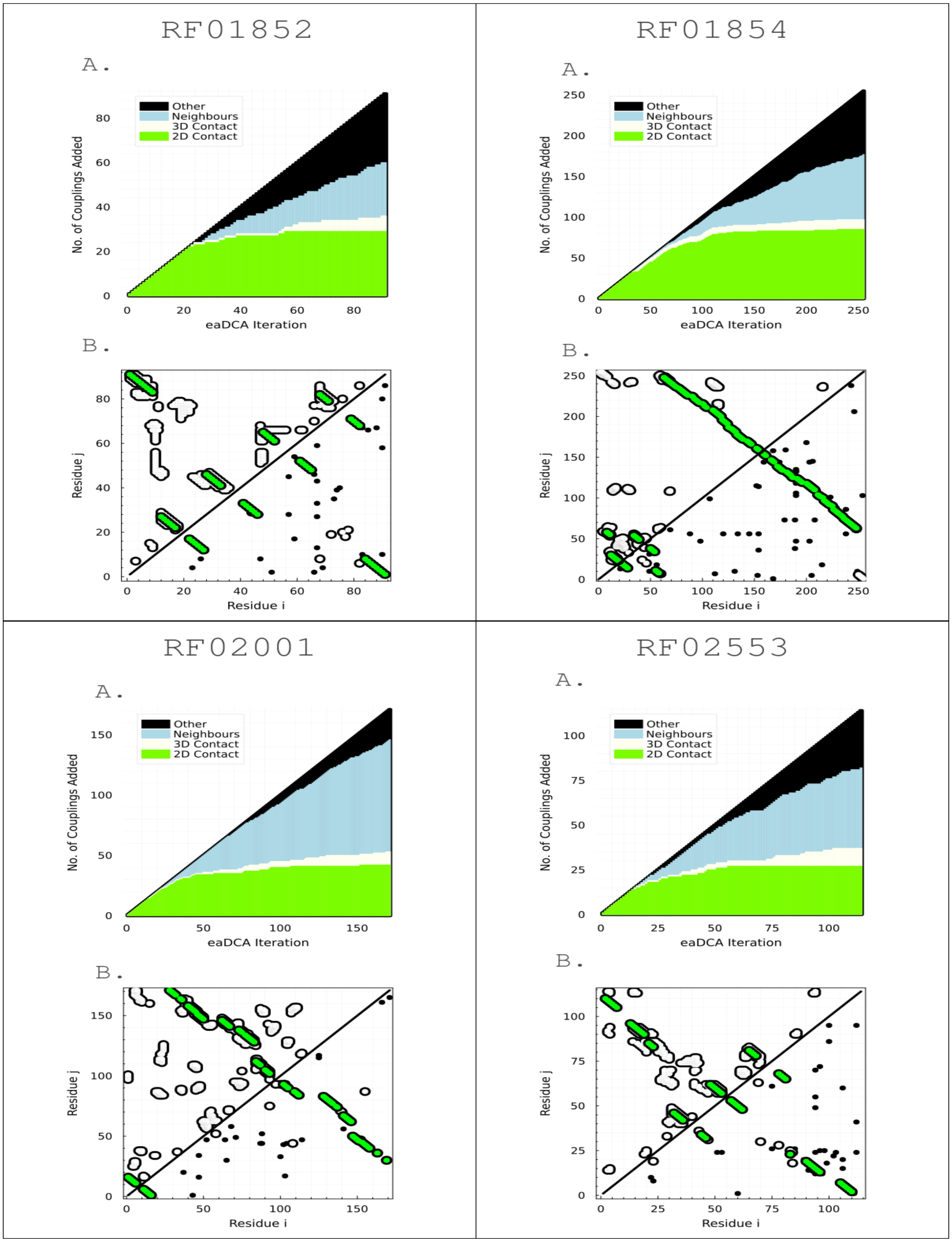


**Figure 3**: **A.** Firs $L$ edges classification. **B.** Contact-Map (upper-left) and added edges (lower-right).

### 4 Prediction of Mutational Effects

#### 4.1 Fitness Dataset Characteristics

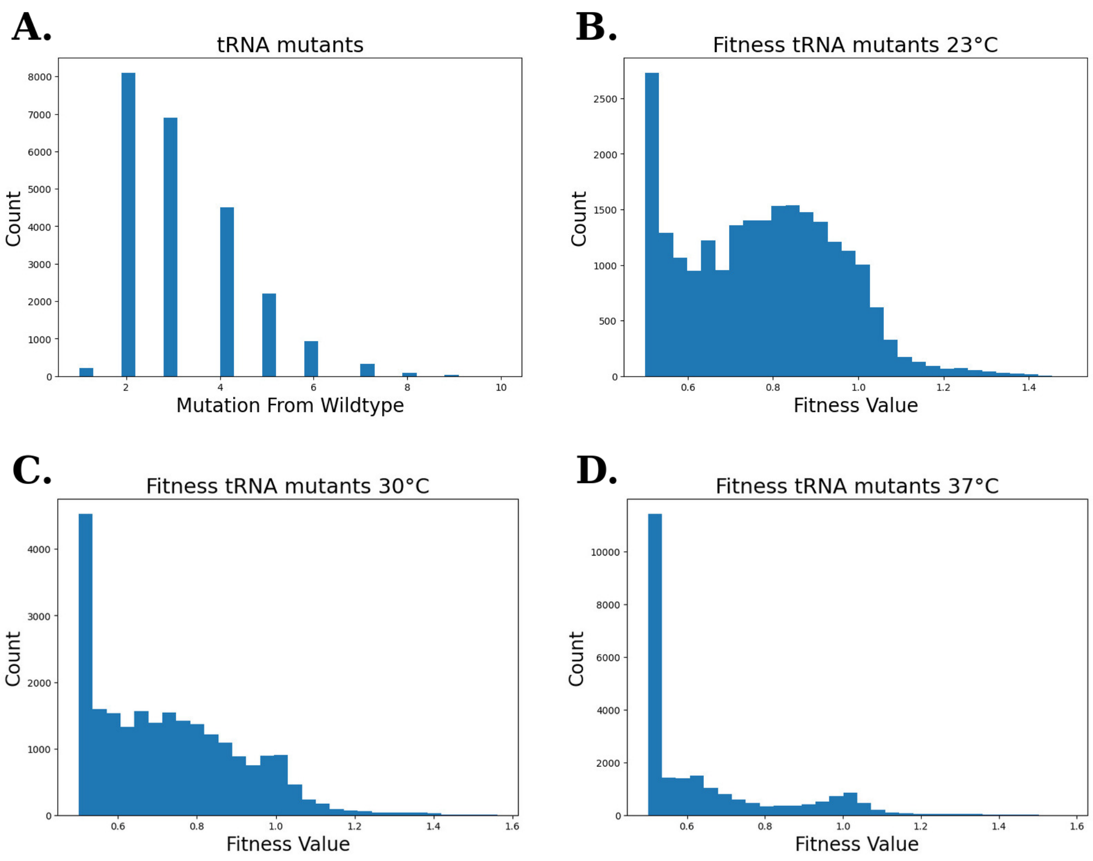


**Figure 4**: **A.** Distribution of the number of mutations from the wildtype. **B.** Fitness distribution at $23°C$. **C.** Fitness distribution at $30°C$. **D.** Fitness distribution at $37°C$.

#### 4.2 Prediction of tRNA Mutational Effects at 23°C, 30°C and 37°C.

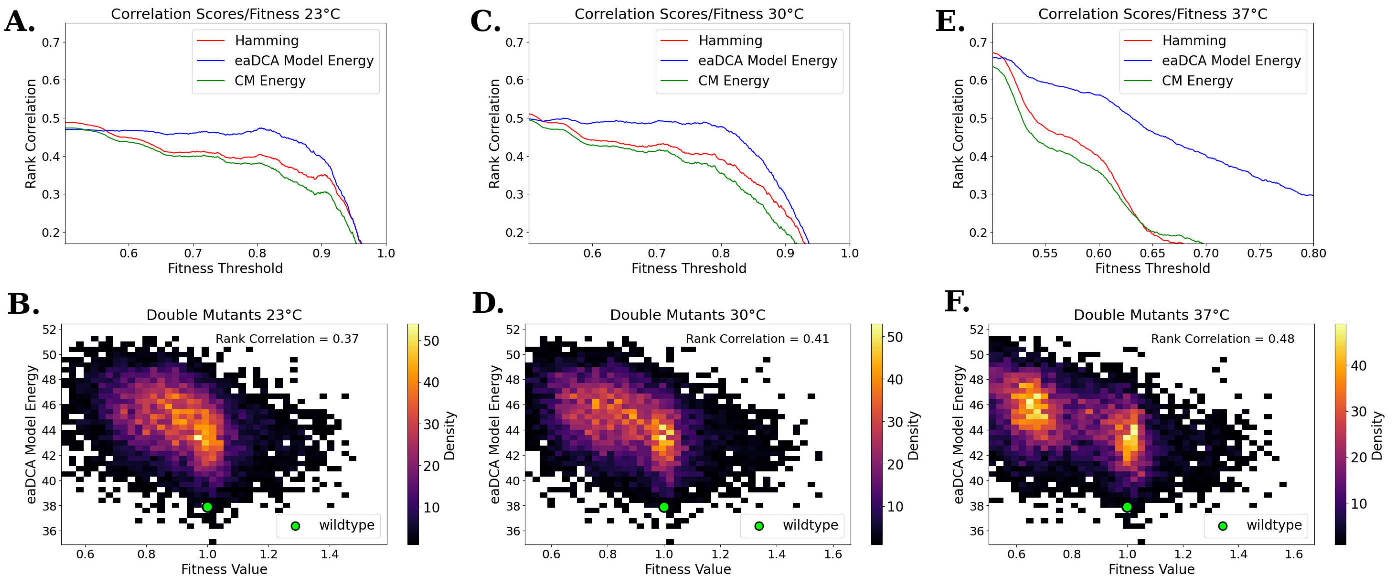


**Figure 5**: **A.** Correlation of Hamming distance, eaDCA model energy and CM energy with tRNA $23°C$ fitness at different values of minimum fitness threshold. **B.** Relation between eaDCA model energy and tRNA $23°C$ fitness on the 8101 double mutants. **C.** Correlation of Hamming distance, eaDCA model energy and CM energy with tRNA $30°C$ fitness at different values of minimum fitness threshold. **D.** Relation between eaDCA model energy and tRNA $30°C$ fitness on the 8101 double mutants. **E.** Correlation of Hamming distance, eaDCA model energy and CM energy with tRNA $37°C$ fitness at different values of minimum fitness threshold (also present in the main text). **F.** Relation between eaDCA model energy and tRNA $37°C$ fitness on the 8101 double mutants (also present in the main text).

### 5 SHAPE-MaP Experiments

#### 5.1 Selection Process For The Tested tRNA

We delineate the two-step selection process employed to select the 76 tested tRNA sequences generated from the eaDCA model.
To begin, 12000 artificial tRNA sequences were generated based on the RF00005 model. During this generation process, we kept the last 16 residues fixed to the yeast tRNA(asp) ones, while also prohibiting the introduction of gap states.
The selection process involved two filtering criteria:

**-** Criteria 1: Secondary Structure

- The secondary structure of yeast tRNA(asp) was predicted using the RNAfold tool from the Vienna RNA (October 2022) package.
- F score $F$ was calculated between the predicted secondary structure and the consensus tRNA secondary structure.
- The pool of 12000 sequences was reduced, retaining only the sequences whose RNAfold-predicted structure had an f-score greater than that of yeast tRNA(asp) $(F > 53)$ .

**-** Criteria 2: eaDCA energy

The secondary structure-filtered dataset was further refined based on the energy of the eaDCA model:

- The 38 sequences of group A were randomly selected among the ones that had an energy lower than that of yeast tRNA(asp) ($E(a_1, \ldots, a_L) < 44$).
- The 38 sequences of group B were randomly selected among the ones that had an energy lower than the 6% left tail of the energy distribution ($E(a_1, \ldots, a_L) < 35$).

This two-step filtering process based on secondary structure and model energy, therefore, resulted in two distinct groups, Group A and Group B, each composed of 38 tRNA sequences.

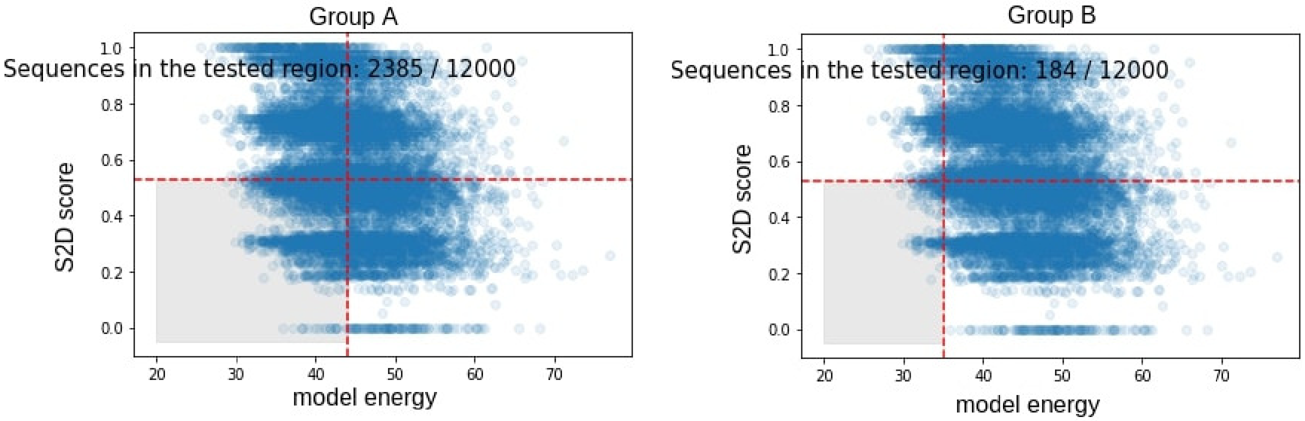


**Figure 6**: To visualize the filtering criterion applied, we shaded in graythe areas from which samples for Group A and Group B were randomly selected.

#### 5.2 Tested tRNA Dataset Characteristics

##### 5.2.1 Sequence Diversity

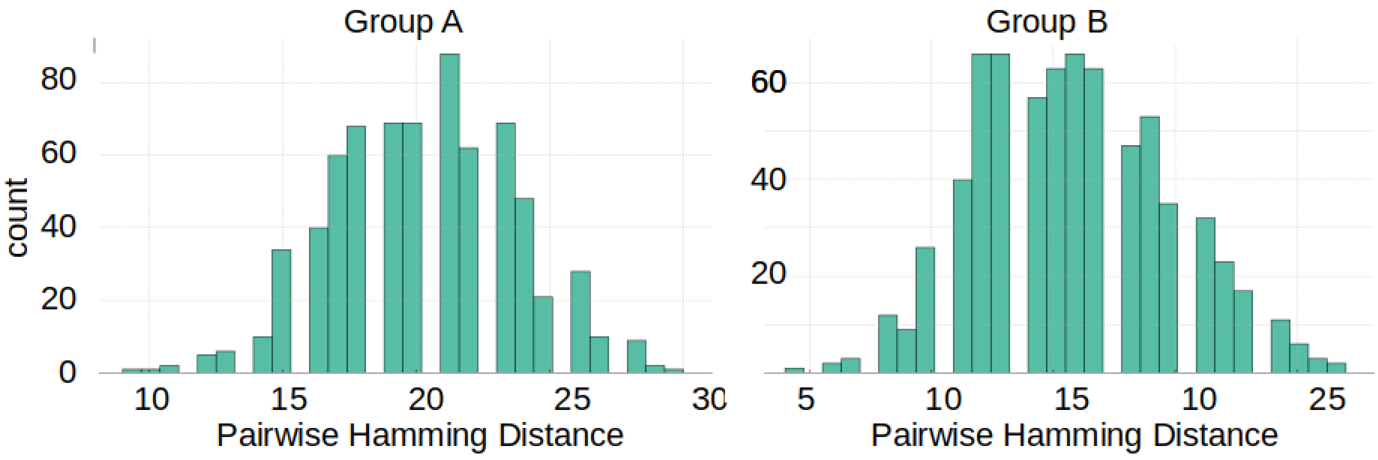


**Figure 7**: Intra-group pairwise sequence distances. This represents the *diversity* of the datasets. Group B, as expected, is less diverse since the energy filtering criterion is more stringent.

##### 5.2.2 Sequence Novelty

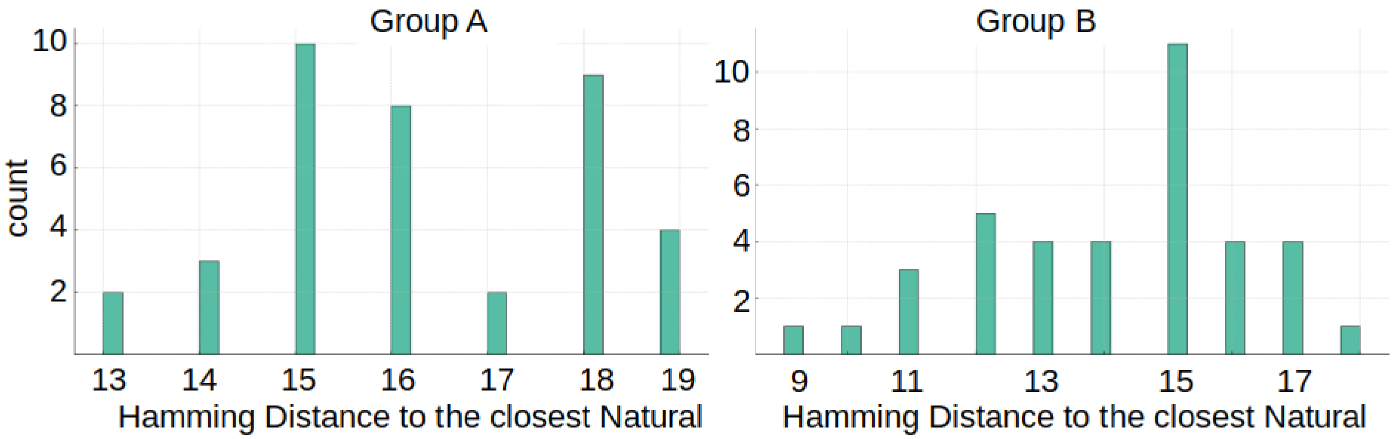


**Figure 8**: Distance from the closest natural for the two groups. This represents the *novelty* of the datasets. Group B, as expected, introduces less novelty since the energy filtering criterion is more stringent. It's important to note that in our generation, the last 16 sites are maintained as constant, matching those of the yeast tRNA-Asp. These sites are not only conserved but also participate in forming secondary structure pairs. Consequently, effectively 32 out of 71 sites are fixed in our investigations. The novel aspects of our study are primarily concentrated on the remaining 39 variable sites.

#### 5.3 SHAPE-MaP Detailed Protocol

##### 5.3.1 RNA Production

The constructs were designed with the T7 promoter positioned at the 5' end of the tRNAs, and with the 15 terminal nucleotides of each tRNA intentionally fixed to facilitate PCR amplification. In total, 77 tRNAs were designed. Out of these, 7 tRNAs were individually produced and modified as single RNA using gBlock DNA templates from Integrated DNA Technologies. The remaining tRNAs were produced and modified as pools of 10 tRNAs using oligoPools from Integrated DNA Technologies.

The DNA templates were amplified by PCR, using 0.5 uM of forward and reverse primers, 0.2 mM of dNTPs (Sigma), 1U of Phusion Hot Start Flex polymerase (New England Biolabs), 1 ng of DNA template, and 1X HF Phusion buffer. The resulting amplified DNA was purified using the NucleoSpin Gel and PCR cleanup kit from Macherey Nagel and quantified using a Nanodrop spectrophotometer. The in vitro transcription was carried in 20 ul reaction volume for 4 h at 37 C, using the HiScribe T7 High Yield RNA Synthesis Kit (New England Biolabs) according to manufacturer instructions, with 800 to 1200 ng of DNA.The transcribed RNAs were then purified by phenol-chloroform extraction using cold acid phenol-chloroform at pH 4.5 (Invitrogen) and ethanol precipitation with 0.1 volume of 3M Sodium Acetate (Sigma) and 2.5 volumes of ice cold 100% ethanol. The precipitated RNAs were resuspended in 50 ul of nuclease-free water and treated with 5U of DNase I (New England Biolabs) for 15 min at 37°C. The RNAs were mixed with loading buffer containing 70% formamide, 130 mM EDTA, 0.1% xylene cyanol, and 0.1% bromophenol blue and loaded onto a pre-run 8% urea polyacrylamide gel electrophoresis (PAGE). The RNAs were then purified from the gel, ethanol precipitated, resuspended in nuclease-free water, quantified using the Qubit RNA Broad Range assay kit (ThermoFisher) and diluted to a concentration of 10 uM.

##### 5.3.2 RNA Modification

5 pmol of RNA in 6 ul of RNase-free water were heated at 95°C for 2 min and placed on ice for 3 min. Subsequently, 3 ul of 3X folding buffer (final concentration: 50 mM Hepes pH 8.0, 200 mM potassium acetate pH 8.0, and 3 mM $MgCl_2$) were added, and the tubes were incubated at 37°C for 20 min to allow the refolding of RNAs. 1 ul of 10X 1M7 (Sigma) in DMSO for the positive reaction (final

concentration : 10 mM 1M7) or 1 ul of neat DMSO for the negative reaction was added and rapidly mixed to ensure even distribution. The solutions were immediately incubated at 37°C on a heating block and the reaction was allowed to proceed for 5 min. Following the modification step, the RNA solutions were cooled on ice and purified by ethanol precipitation. For the denaturing condition, 5 pmol of RNA in 3 ul in RNase-free water were mixed with 5 ul of 100% highly deionized formamide (Hi-Di Formamide, Applied Biosystems) and 1 ul of 10X denaturation buffer (final concentration : 50 mM Hepes pH 8.0, 40 mM EDTA). The tubes were incubated at 95°C for 1 min to denature the RNAs. Similarly, 1 ul of 10X SHAPE reagent in DMSO was added to the denatured RNAs, rapidly mixed, and immediately incubated at 37°C for 5 min. After the modification step, the solutions were cooled on ice and purified alongside the negative and positive reactions.

The ethanol precipitation was performed as follows: the 10 ul of modified RNAs were diluted with 87 ul of RNase-free water, and mixed with 1 ul of 20 mg/ml glycogen (Invitrogen) and 2 ul of 100 mM EDTA pH 8. Next, 0.1 volume of 3 M NaAc and 3.5 volumes of ice cold 100% ethanol were added to the solution, and the RNas were precipitated overnight at -20°C, centrifuged at 13,000 rpm for 1 h at 4°C, and resuspended in 15 ul of RNase-free water. The RNA concentrations were quantified using the Qubit RNA High Sensitivity assay kit (ThermoFisher).

##### 5.3.3 Library Preparation

After purification, the modified RNAs were pooled in equimolar proportions based on their conditions (positive, negative, and denaturing) for reverse transcription. The reverse-transcription was carried out with 1 pmol of RNA in 20 ul of reaction volume, as follows : i) the RNAs were incubated at 95°C for 3 min and transferred to ice, ii) 1 ul of a 2 uM reverse primer, complementary to the end of the tRNA and carrying the Rd2 adaptor, was added to the solutions and incubated at 65°C for 5 minutes and transferred to ice, iii) 8 ul of freshly prepared 2.5X Shape-Map buffer (final concentration: 50 mM Tris-HCl pH 8.3, 75 mM KCl, 10 mM DTT, 6 mM MnCl2, 0.5 mM dNTP mix) was added and incubated for 2 min at 50°C before being placed on ice, iv) 200 U of SuperScript II reverse transcriptase (ThermoFisher) and 1 ul of 100 uM Rd1-TSO were added and incubated at 42°C for 3 hours, followed by an inactivation step at 70°C for 15 minutes. The resulting cDNAs were purified with AMPure XP beads (Beckman Coulter) with a bead-to-sample ratio of 1.8, and eluted in 15 ul of RNase-free water.

A PCR enrichment was carried out for 8 cycles, using 25 ul of KAPA HiFi HotStart ReadyMix (Roche), 5 ul of each primer (NEBNext Multiplex Oligos for Illumina, New England Biolabs), and 15 ul of purified cDNAs. The DNA libraries were purified with AMPure XP beads (Beckman Coulter) with a bead-to-sample ratio of 0.9, and eluted in 20 ul of RNase-free water, and quantified using the Qubit dsDNA High Sensitivity assay kit (ThermoFisher) and quantitative PCR (KAPA Library Quantification Kit, Roche). The length distribution of the libraries was analyzed using the High Sensitivity D1000 ScreenTape for TapeStation Systems (Agilent). The libraries were sequenced on a MiSeq-V3 flow cell (Illumina) in paired ends with 25% of PhiX by the NGS platform at Institut Curie (Paris, France).

| **Primer Name** | **Sequence (5'-3')** |
|---|---|
| tRNA_R | TGCCGCGACGGGGAA |
| tRNA_F | TAATACGACTCACTATAG |
| TSO_Rd1 | TACACGACGCTCTTCCGATCTrGrGrG |
| Rd2_tRNA | GTGACTGGAGTTCAGACGTGTGCTCTTCCGA-TCTTGCCGCGACGGGGAA |
| i5 primer | AATGATACGGCGACCACCGAGATCTACACTC-TTTCCCTACACGACGCTCTTCCGATC-s-T |
| i7 primer | CAAGCAGAAGACGGCATACGAGAT-NNNNNN-GTGACTGGAGTTCAGACGTGTGCTCTTCCGATC-s-T |

**Table 3**: Sequences of the primers used. -s- indicates a phosphorothioate bond. N indicates random nucleotide

##### 5.3.4 SHAPE Reactivity Profiles

Here we show the projection of SHAPE reactivity onto the tRNA consensus secondary structure for tested sequences (and for another natural tRNA took from the SHAPE reference dataset). There doesn't appear to be a significant difference between groups A and B among the sequences that exhibit a compatible profile.

01_A CRITERIA MET

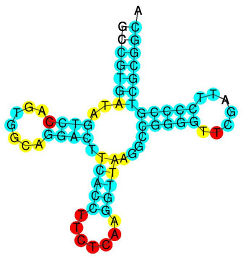

03_A

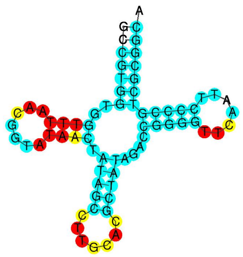

03_B

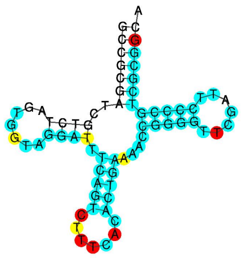

04_B

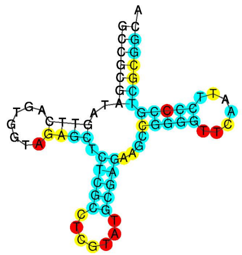

05_A

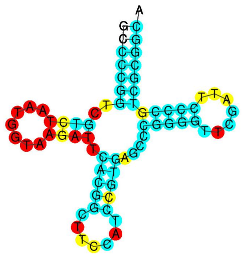

07_A

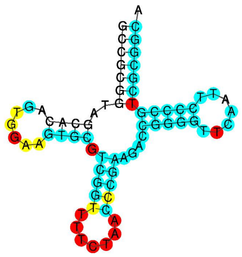

08_A CRITERIA MET

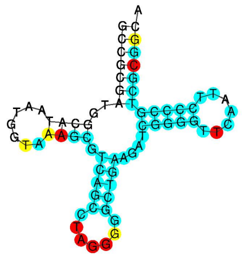

10_A CRITERIA MET

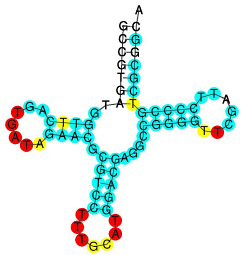

10_B CRITERIA MET

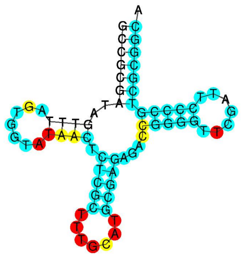

12_B

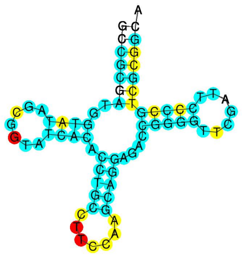

13_A

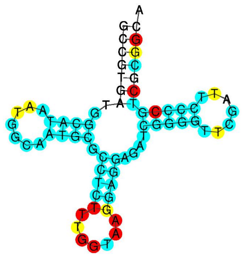

13_B CRITERIA MET

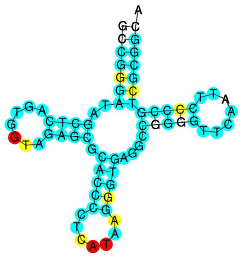

16_A CRITERIA MET

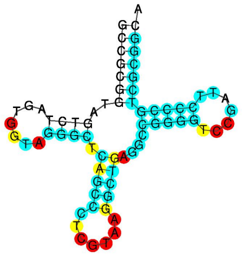

16_B CRITERIA MET

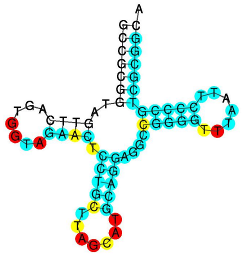

18_A

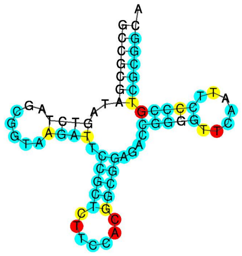

19_A CRITERIA MET

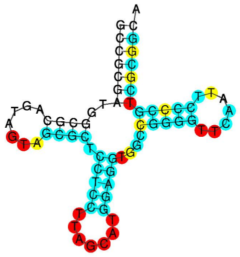

20_A CRITERIA MET

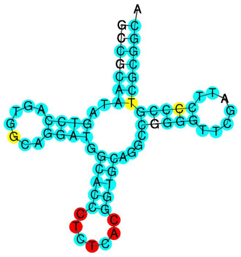

21_B CRITERIA MET

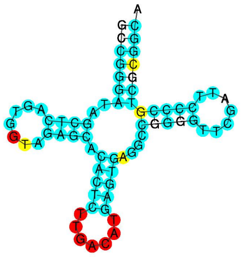

22_A

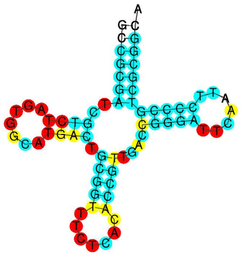

23_A

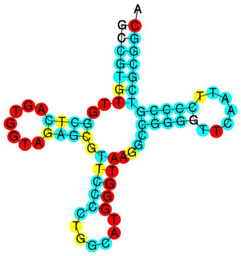

24_B

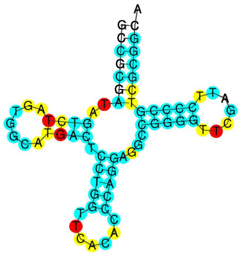

25_B CRITERIA MET

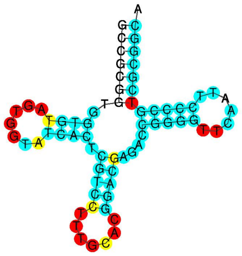

26_A

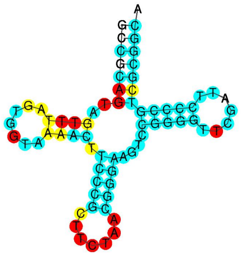

27_B

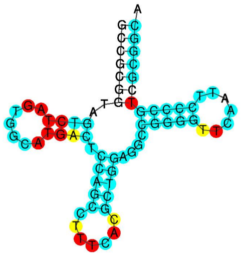

29_A

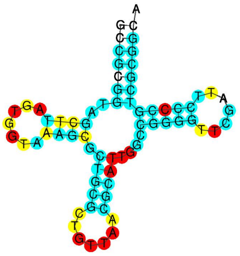

30_A

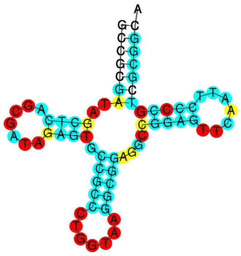

31_A

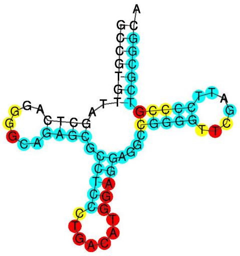

31_B

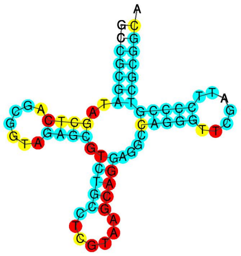

32_A

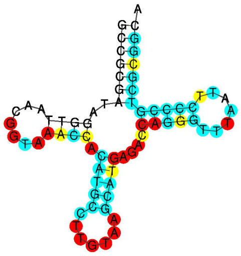

33_A

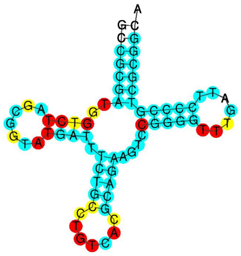

34_A CRITERIA MET

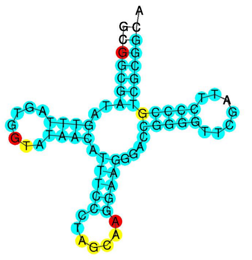

37_A

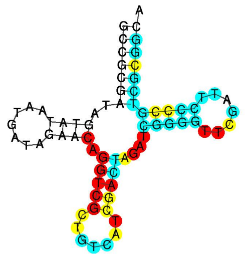

38_A

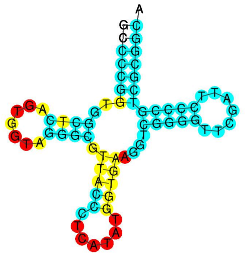

38_B CRITERIA MET

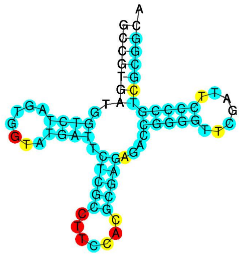

tRNA_PHE_E_COLI

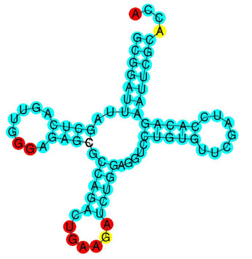
